# The Wnt1-Cre2 transgene causes aberrant recombination in non-neural crest cell types

**DOI:** 10.1101/2024.11.06.622365

**Authors:** Shashank Gandhi, Emily J. Du, Edivinia S. Pangilinan, Richard M. Harland

## Abstract

The Wnt1-Cre2 driver, designed to address the effect of Wnt1 overactivation in the ventral neural tube in the original Wnt1-Cre line, was recently shown to have ectopic expression in the male germline. When crossed with a reporter mouse, we observed fluorescent protein expression in non-neural-crest cell types in the gut. Here, we characterize the pattern of Cre-mediated recombination in the Wnt1-Cre2 driver using three transgenic reporter lines. We find aberrant reporter activation in the gut endoderm in embryonic and postnatal timepoints, starting as early as E8.5. This pattern of recombination was independent of the age, sex, and type of reporter line used, with the Wnt1-Cre2 allele inherited from either sires or dams resulting in ectopic fluorescence in the intestinal epithelium. We also detect reporter activity in the ventral neural tube. However, expression in the neural crest and its derivatives remained consistent with previous studies. We further quantify differences in the non-specific recombination observed across reporter lines using flow cytometry. Interestingly, the penetrance of reporter activation between reporter lines was different, with R26R^*mTmG*^ showing less ectopic activation than the R26R^*tdTom*^ and R26R^*eYFP*^ lines. Finally, we propose a potential mechanism whereby genes surrounding the Wnt1-Cre2 insertion site on mouse chromosome 2 contribute to its Wnt1-independent activation in the endoderm. Taken together, our results suggest that users should exercise caution when using the Wnt1-Cre2 driver line for neural crest studies in the mouse.

## Introduction

The neural crest is a multipotent population of cells that originates at the neural plate border and migrates extensively during embryogenesis (Le Douarin, 1982). Historically, lineage tracing of neural crest derivatives in amniotes used quail-chick chimeras generated by transplantation of dorsal neural tubes from quail embryos into chick hosts (Bronner and Simões-Costa, 2016; Gandhi and Bronner, 2021; Le Lièvre and Le Douarin, 1975). However, genetically encoded tracers in mice enable sophisticated and reliable lineage tracing (Tang and Bronner, 2020; Yoshida et al., 2008). The now ubiquitous Cre/lox system (Sauer and Henderson, 1988; Sternberg and Hamilton, 1981), for example, enables the indelible labeling of cells and their derivatives in addition to lineage-specific gene deletions for functional characterization (Rossant and Nagy, 1995).

The *Wnt1* gene is expressed in the dorsal neural tube (McMahon and Bradley, 1990; Thomas and Capecchi, 1990) in progenitors of the neural crest lineage, and its promoter was used in mice to drive Cre recombinase expression in the neural crest (Danielian et al., 1998). Recombinase activation of lineage tracers confirmed neural crest contributions to mammalian development, and the line was further used for conditional deletion of genes to generate animal models of neural-crest-associated congenital defects, including Treacher Collins syndrome (Trainor, 2010), CHARGE syndrome (Sperry et al., 2014), and Hirschsprung Disease (Bondurand and Southard-Smith, 2016). However, the Wnt1-Cre transgene also expressed additional *Wnt1* in the midbrain, resulting in differentiation defects of ventral midbrain neurons. To circumvent these aberrant phenotypes, the Wnt1-Cre2 driver line was generated with the same regulatory elements but without the 5’ UTR segment of the Wnt1 gene which was suspected to cause the overexpression phenotypes (Lewis et al., 2013).

The Wnt1-Cre2 driver line has recently been reported to activate Cre recombinase ectopically. Dinsmore and Soriano (Dinsmore et al., 2022) found germline activation of the Cre gene associated with its expression in the spermatogonia of adult male mice. Moreover, additional anecdotes suggested inappropriate activation of the Wnt1-Cre2 driver (Drokhlyansky et al., 2020). While using this transgenic line to study gut development, we also found ectopic sites of activation, leading us to analyze this phenomenon further. In this paper, we address the effect and potential mechanism of ectopic activation using the R26R^*tdTom*^ (Madisen et al., 2010), R26R^*eYFP*^ (Srinivas et al., 2001), and R26R^*mTmG*^ (Muzumdar et al., 2007) reporter lines. We found unexpected reporter activity in the gut endoderm following Wnt1-Cre2-mediated recombination in embryos and postnatal pups of the C57BL/6J strain, with varying levels of penetrance. We then used flow cytometry to quantify the extent of Cre activation in the endoderm. Finally, we have explored possible reasons for this phenomenon by analyzing an existing single-cell RNA-sequencing dataset from the mouse small intestinal epithelium (Han et al., 2020).

Taken together, our findings underscore the need for caution and orthogonal validation approaches when using the Wnt1-Cre2 driver line for neural crest studies in the mouse.

## Methods

### Mouse husbandry

The Animal Use Protocol under which all animal experiments were conducted was approved by the Institutional Animal Care and Use Committee of the University of California, Berkeley. All mice were housed on a 14h-10h light-dark cycle with access to food and water *ad libitum* in an animal vivarium monitored daily by dedicated in-house technicians and veterinary staff of the Institutional Office of Lab Animal Care.

### Mouse strains and dissections

The following strains were used in this study:

| Strain | Common name | Source | Identifier |
| --- | --- | --- | --- |
| C57BL/6J | B6 | JAX | 000664 |
| B6.Cg-Gt(ROSA)26Sortm14(CAG-tdTomato)Hze/J | R26R<sup>tdTom</sup> | JAX | 007914 |
| B6.Cg-Tg(Wnt1-cre)2Sor/J | Wnt1-Cre2 | JAX | 022501 |
| B6.129x1-Gt(ROSA)26Sortm1(EYFP)Cos/J | R26R<sup>eYFP</sup> | JAX | 006148 |
| B6.129(Cg)-Gt(ROSA)26Sortm4(ACTB-tdTomato,-EGFP)Luo/J | R26R<sup>mTmG</sup> | JAX | 007676 |
| Wnt1-Cre2<sup>Tg/+</sup> ; R26R<sup>tdTom/+</sup> | Wnt1-Cre2-tdTom | In-house breeding | - |

All mice used in this study were maintained on a C57BL/6J background. Timed matings were set up an hour before the beginning of the dark cycle. The next morning, the dam was checked for a vaginal plug and annotated as E0.5 if the plug was present. All embryos presented in this study were dissected in 1x PBS and fixed in 4% paraformaldehyde (PFA) overnight at 4°C.

The intestines from postnatal pups were dissected and transferred to 1x PBS. Small pieces were then selected and cut using epifluorescence microscopy, fixed overnight at 4°C in 4% PFA, and embedded for sectioning as described below.

### Cryosectioning and Immunohistochemistry

The tissue was cryopreserved in 15% sucrose and embedded in OCT. Blocks were stored at -80°C until they were sectioned. 10-14μm sections were collected on charged microscope slides and stored overnight at room temperature, and long-term at -20°C.

All reporters besides R26R^*eYFP*^ were bright enough to be imaged directly, but R26R^*eYFP*^ samples were immunostained for GFP. Briefly, slides were equilibrated to room temperature and soaked in 1x PBS in a Coplin jar in a 37°C water bath to dissolve the OCT. Following two additional washes in PBS, the slides were blotted dry and blocked in 10% donkey serum in 0.1% PBS-Triton for 1 hour at room temperature. A primary antibody against GFP (Rockland technologies, AB_218182, 1:500) was dissolved in the blocking solution and the slides were incubated overnight at 4°C. The next day, the slides were washed in 1x PBS four times for fifteen minutes each. The secondary antibody (Donkey anti-goat Alexa Fluor 488; 1:500) was dissolved in the blocking solution and incubated on the sections for 1 hour at room temperature. We also added DAPI (1:5000) to the secondary antibody solution. The slides were washed twice in 1x PBS for fifteen minutes each, followed by one wash in deionized water. Sections were air-dried and mounted with a glass coverslip using Fluoromount-G. Slides were left overnight before imaging.

### Genotyping

All embryos were genotyped using protocols recommended by the Jackson Laboratory. A gradient PCR was first set up for each primer pair to check the optimal annealing temperature on a positive control animal. The following primer pairs were used for genotyping animals in this study:

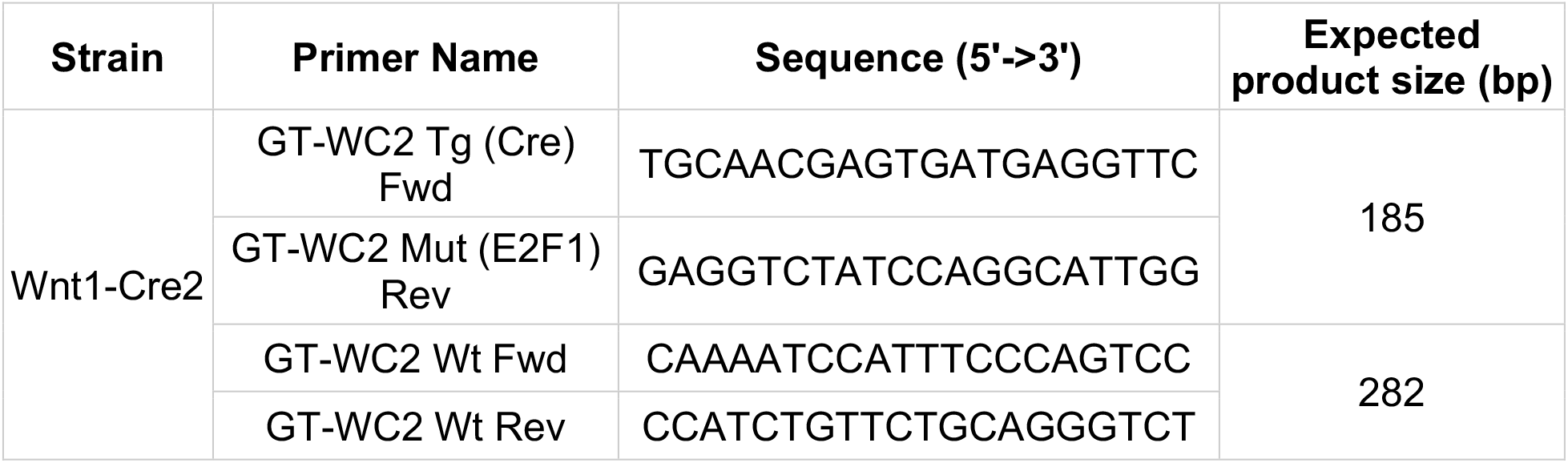
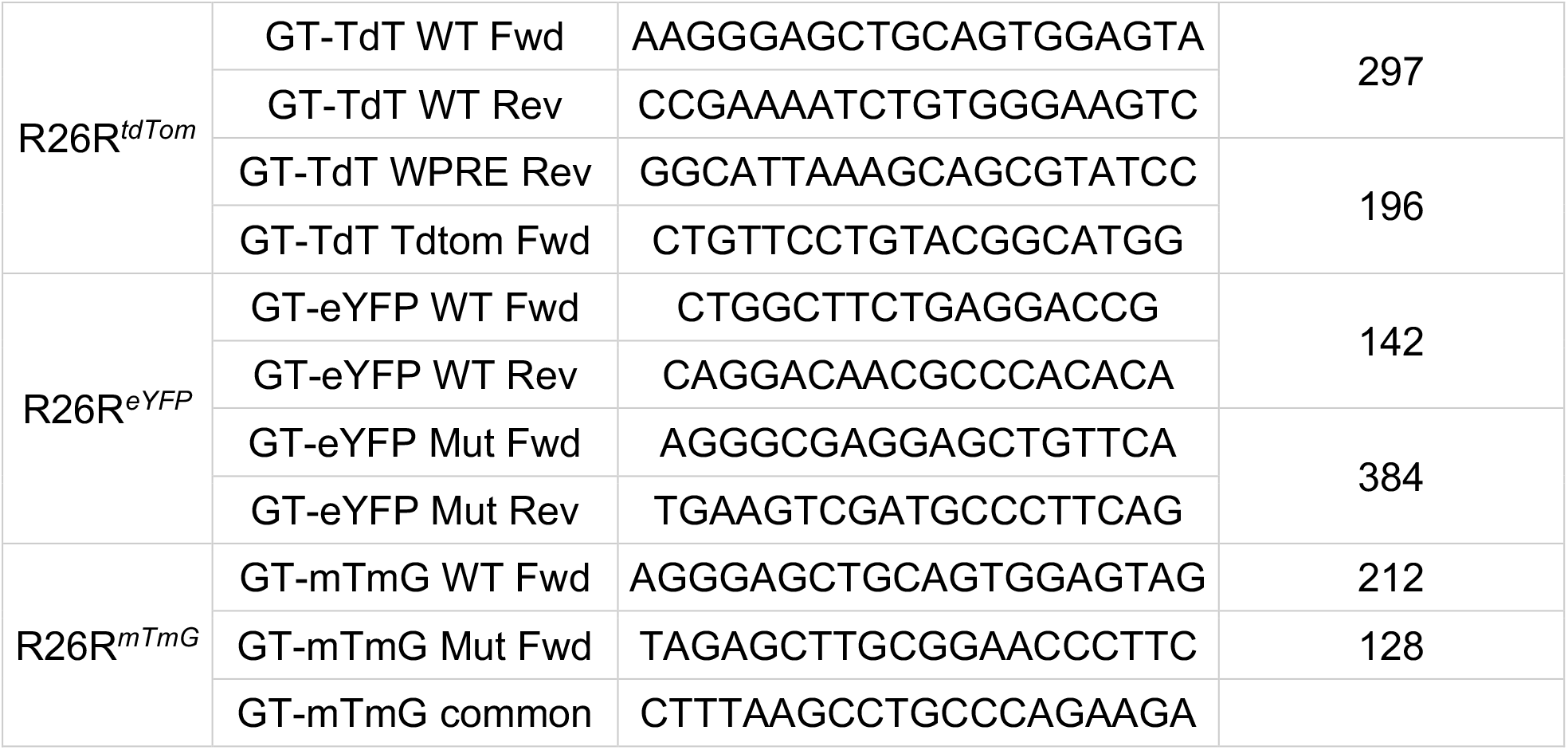

### Epifluorescence and confocal microscopy

Wholemount images were taken with a Leica Thunder fluorescence dissecting microscope, whereas sections were imaged on a Zeiss LSM980 upright confocal or a Zeiss LSM900 inverted confocal. Z-stacks were collected using the optimal step size recommended for each objective, and maximum intensity projections were generated in Zen Blue (Zeiss). Level adjustments and LUT allocations were made in ImageJ (Schindelin et al., 2012).

### Flow cytometry

E15.5 Wnt1-Cre2^*Tg/+*^; R26R^*eYFP/+*^ and Wnt1-Cre2^*Tg/+*^; R26R^*mTmG/+*^ embryos were dissected in 1x PBS. Intestines were isolated from the body cavity and carefully unfolded to remove mesentery whilst preserving the muscularis. The midgut of each embryo was macerated with scissors in an individual 1.7mL Eppendorf tube and washed once with 1x PBS. Enzymatic dissociation was carried out with 0.5% Trypsin-EDTA at 37°C for approximately 25 minutes, with mechanical trituration every 5 minutes. Samples were then filtered through a 40μm cell strainer into Hank’s-0.5% BSA and centrifuged at 500g for 5 minutes at 4°C. Samples were resuspended in 250μL of red cell lysis buffer (Roche) at room temperature for one minute, washed in cold 1x PBS, and stained on ice in the dark with LIVE/DEAD Blue (Life Technologies; 1:200) in 1x PBS in a final volume of 100μL for 30 minutes. Following one wash in 1x PBS and 2 washes in Hanks-0.5% BSA, cells were resuspended in Rat IgG2a anti-mouse EpCAM conjugated to BV421 (Biolegend, AB_2563983; 1:200) in Hanks-0.5% BSA and stained on ice in the dark for 50 minutes. Following three washes in Hanks-0.5% BSA, cells were transferred to FACS tubes and analyzed on a 5-laser Cytek® Aurora. Reference controls were generated with stage-matched wild-type E15.5 embryonic gut tissue. An isotype control Rat IgG2a antibody conjugated to BV421 (Biolegend, AB_10933427; 1:200) was used to validate the EpCAM staining. All data was analyzed in FlowJo™ (BD) and plots assembled in Inkscape.

### Single-cell RNA-sequencing analysis

To characterize the potential gut expression of genes neighboring the Wnt1 transgene, we reanalyzed expression counts from Han et al., 2020. The count matrix associated with GSE136689 (NCBI GEO) was imported in R for analysis using the Seurat library (Butler et al., 2018). Briefly, cells with more than 500 UMIs, less than 20,000 RNA counts, and less than 15% mitochondrial gene expression were selected for downstream analysis. The first 15 principal components were used to reduce the dimensionality of the data. The data split into two main clusters: definitive endoderm (DE) and splanchnic mesoderm (SM). DE cells were extracted from this data and *Epcam* expression was used to identify 4000 epithelial cells. Using the metadata provided by Han et al., 2020, each cell was labeled with its associated embryonic time point. This information was used to regress this variable from the data. To identify genes neighboring the *E2f1* locus on the 2nd chromosome where the Wnt1-Cre2 transgene is inserted, we selected all protein-coding genes with any part of their coding sequence within a 300kb region surrounding *E2f1*. Finally, gene expression was visualized using built-in Seurat functions *FeaturePlot, DimPlot*, and *RidgePlot*.

### Statistics

To compare the recombination frequency in Wnt1-Cre2^*Tg/+*^; R26R^*eYFP/+*^ and Wnt1-Cre2^*Tg/+*^; R26R^*mTmG/+*^ embryos, we used a two-tailed unpaired t-test in R. Significance was calculated based on a p-value cutoff of 0.05. Histograms and dot plots were plotted in R using the ggplot2 package. All plots were assembled into figures in Inkscape.

## Results and Discussion

We were originally interested in using the Wnt1-Cre2 driver line to label neural-crest-derived enteric neurons in the adult mouse gut. To that end, we obtained Wnt1-Cre2^*Tg/+*^ mice from JAX and crossed them with the R26R^*tdTom*^ reporter line to obtain Wnt1-Cre2^*Tg/+*^; R26R^*tdTom/+*^ pups. We dissected a P5 male pup to isolate the intestine and validate neural crest labeling in the gut. However, upon dissection, we observed patches of strong tdTomato signal as bright puncta along the length of the digestive tract [Figure 1A-B], together with signal in the neural-crest derived enteric neurons [Figure 1C]. To further investigate the origin of the non-specific signal, we dissected different segments of the gut and sectioned them. We observed long strips of Cre-mediated reporter activation along the length of the villus, originating from the intestinal crypt [Figure 1D-E’]. This was reminiscent of “ribbons” [Figure 1E] spanning the entire crypt-villus axis reported in seminal lineage tracing experiments of the stem cells residing in the crypt (Snippert et al., 2010). To make sure that the recombination phenotype was not sex-specific, we also dissected a female pup which [Figure 1F-F’’] demonstrated similar levels of recombination in the villi. Moreover, when the genotypes of the dam and sire were reversed, such that Wnt1-Cre2^*Tg/+*^ females were crossed with R26R^*tdTom*^ homozygous males, the outcome was similar. Overall, we analyzed 8 Wnt1-Cre2^*Tg/+*^; R26R^*tdTom/+*^ pups (the Wnt1-Cre2 allele was inherited by 6 pups from the dam and by 2 pups from the sire) across 5 different litters, with 100% of the samples exhibiting non-neural crest activation in the endoderm [Table 1].

**Table 1:** Endoderm recombination in postnatal pups.

| Paternal Genotype | Maternal Genotype | Total pups dissected | Endoderm recombination frequency (%) |
| --- | --- | --- | --- |
| R26R<sup>tdTom/tdTom</sup> | Wnt1-Cre2<sup>Tg/+</sup> | 6 | 6 (100) |
| Wnt1-Cre2<sup>Tg/Tg</sup> | R26R<sup>tdTom/tdTom</sup> | 2 | 2 (100) |
| R26R<sup>mTmG/mTmG</sup> | Wnt1-Cre2<sup>Tg/+</sup> | 6 | 6 (100) |
| Wnt1-Cre2<sup>Tg/Tg</sup> | R26R<sup>mTmG/mTmG</sup> | 9 | 9 (100) |
| R26R<sup>eYFP/eYFP</sup> | Wnt1-Cre2<sup>Tg/+</sup> | 3 | 3 (100) |
| Wnt1-Cre2<sup>Tg/Tg</sup> | R26R<sup>eYFP/eYFP</sup> | 5 | 5 (100) |

**Figure 1:**
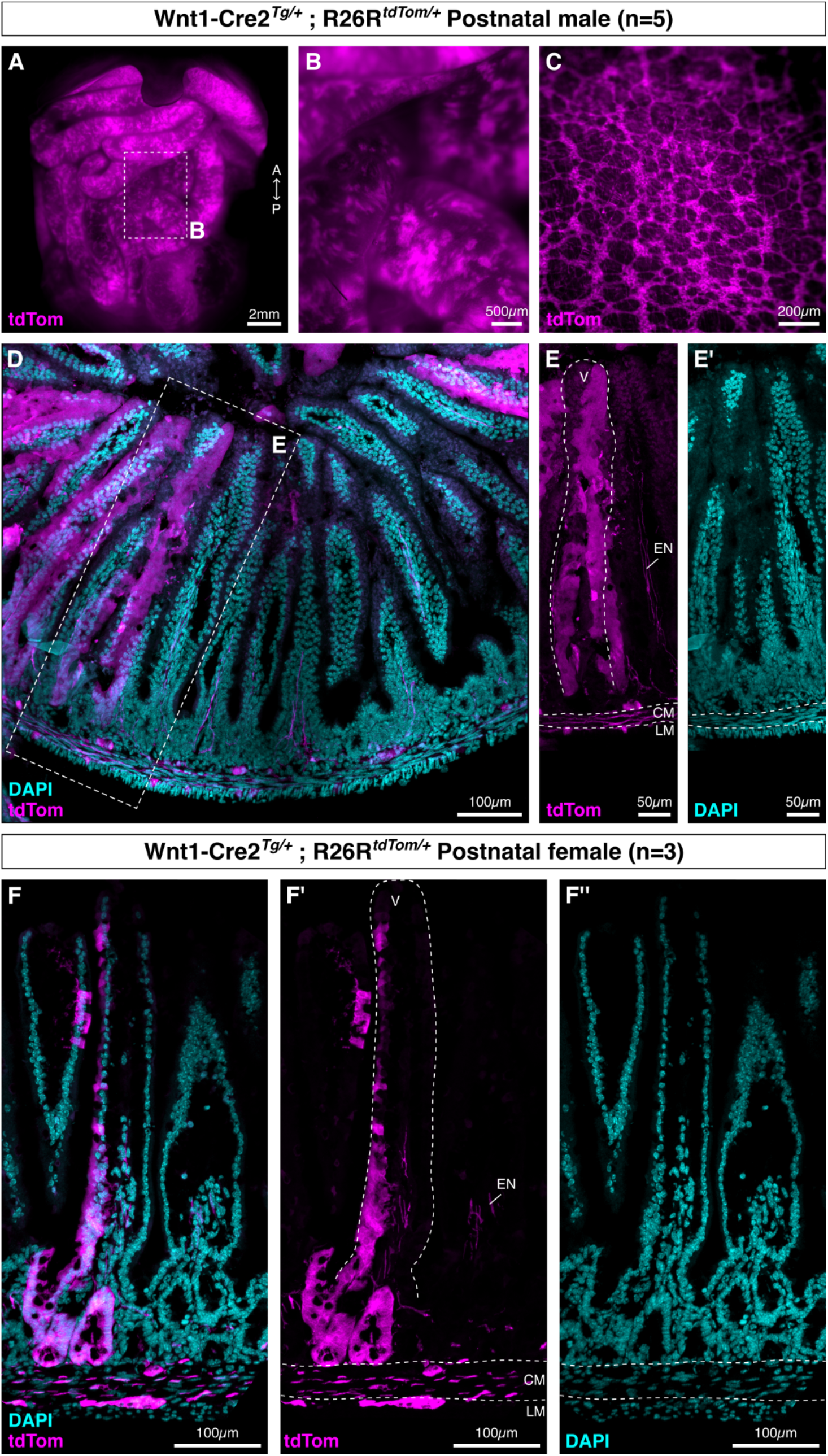
Wnt1-Cre2 transgene drives reporter recombination in the postnatal gut. **A**. A dissected postnatal day (P) 15 male Wnt1-Cre2^*Tg/+*^; R26R^*tdTom/+*^ pup. **B-C**. Zoomed-in views of the intestine shown in A. **D**. A transverse section through the P15 male small intestine. **E-E’**. Reporter activity (E) and nuclei (E’) in the villi and *muscularis* layers of a P15 small intestine. **F**. Transverse section through the small intestine of a P25 female Wnt1-Cre2^*Tg/+*^; R26R^*tdTom/+*^ pup. **F’-F’’**. Reporter activity (F’) and nuclei (F’’) in the villus and muscularis layers of the P25 female small intestine. (Abbreviations: A, Anterior; P, Posterior; EN, Enteric Neurons; CM, Circular Muscle; LM, Longitudinal Muscle; V, Villus)

We reasoned that the aberrant recombination could be a result of either spontaneous activation of Cre recombinase in the crypt stem cells, or carryover of reporter signal from the embryonic endoderm. To explore this further, we set up crosses to obtain a time course of Wnt1-Cre2^*Tg/+*^; R26R^*tdTom/+*^ embryos ranging from E8.5, when neural crest cells from the cranial axial level have delaminated and migrated laterally (Yoshida et al., 2008), to E15.5, when migration of cardiac and vagal neural crest cells into the heart and gut, respectively, has completed (George et al., 2020). At E8.5, we not only observed mosaic activation of the reporter [Figure 2A-B] in the neural crest but also segments of the definitive endoderm that were labeled (n=17). Cross-sections through these embryos [Figure 2C] revealed strong signal within the branchial arch mesenchyme [Figure 2D], which is derived from the cranial neural crest (Chai et al., 2000). However, the reporter was also active within the epithelial cells of the developing gut tube [Figure 2E]. Endodermal signal persisted in E9.5 embryos, where we observed reporter signal in the distal part of the gut tube in wholemount embryos [Figure 2F-F’], even though vagal neural crest cells at the E9.5 stage have not migrated that far (Hutchins et al., 2018).

**Figure 2:**
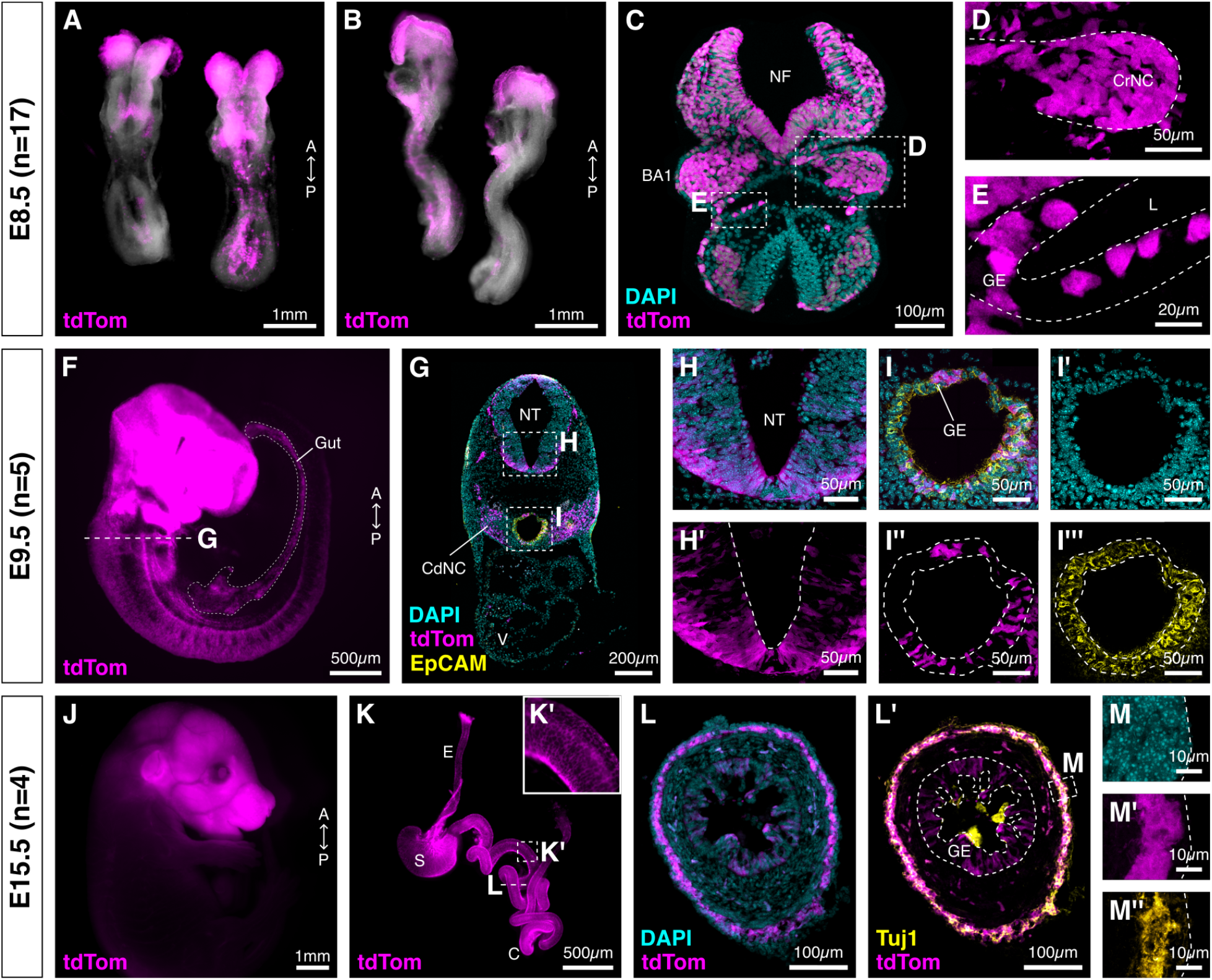
Wnt1-Cre2 transgene drives reporter recombination in the neural crest and ectopically in the gut. **A-B**. Dorsal (A) and Ventral (B) view of two E8.5 Wnt1-Cre2^*Tg/+*^; R26R^*tdTom/+*^ littermate embryos. **C**. A transverse cross-section through a representative E8.5 embryo. **D-E**. Magnified view of the first branchial arch (D) and the developing gut epithelium (E) in E8.5 embryos boxed in C. **F**. Lateral view of an E9.5 Wnt1-Cre2^*Tg/+*^; R26R^*tdTom/+*^ embryo. The developing gut tube is labeled with a dotted line. **G**. Transverse cross-section through the 3^rd^ branchial arch of the embryo shown in F. **H-H’**. The ventral neural tube at E9.5. **I-I’’’**. The gut epithelium at E9.5. I’’ shows ectopic recombination in the EpCAM positive cells (I’’’). **J**. Lateral view of an E15.5 Wnt1-Cre2^*Tg/+*^; R26R^*tdTom/+*^ embryo. **K-K’**. Wholemount view of the digestive tract dissected from embryo shown in K. Neuronal fibers derived from the vagal neural crest are visible (K’). **L-L’**. A transverse section through the small intestine of the embryo shown in K. Enteric neurons (L’) are labeled with Tuj1. **M-M’’**. Magnified view of the myenteric plexus of the embryo shown in K shows neural-crest-derived enteric neurons (M’) stained with DAPI (M) and Tuj1 (M’’). (Abbreviations: A, Anterior; P, Posterior; NF, Neural Folds; BA, Branchial Arch; CrNC, Cranial Neural Crest; GE, Gut Epithelium; L, Lumen; NT, Neural Tube; CdNC, Cardiac Neural Crest; V, Ventricle; E, Esophagus; S, Stomach; C, Caecum)

Upon sectioning these embryos [Figure 2G], we made two notable observations. First, the ventral neural tube along the entire body axis was labeled [Figure 2H-H’]. This was interesting because the Wnt1-Cre2 driver line was generated in response to concerns about ectopic activation of the Wnt1 transgene in the ventral midbrain (Lewis et al., 2013), which caused an overexpression of Wnt1 in the ventral neural tube resulting in increased cell proliferation and loss of important neuronal subtypes, including the dopaminergic neurons. While we observed reporter activity in the ventral neural tube [Figure 2H-H’], we did not check if the reporter activity correlated with increased Wnt signaling, or whether the rate of cell proliferation was affected. Second, we observed that only a fraction of the epithelial cells, but not all, were labeled with tdTomato [Figure 2I-I’’’], suggesting that Cre activity is either stochastic or restricted to a specific sub-population of the developing endoderm.

Finally, we confirmed the presence of vagal neural crest-derived enteric neurons in the E15.5 embryonic gut [Figure 2J]. We examined the digestive tract of these embryos for reporter activity in wholemount [Figure 2K-K’] and cross-sections. As expected, the reporter activity was abundant within the gut epithelium [Figure 2L] and the developing plexuses, which we validated by looking at reporter activity overlap with the expression of the neuronal marker Tuj1 [Figure 2L’-M’’]. Together, these results confirm that the Wnt1-Cre2 transgene robustly mediates reporter expression in non-neural crest cell types in the C57BL/6J strain [Table 2].

**Table 2:** Endoderm recombination in embryos.

| Paternal Genotype | Maternal Genotype | Total embryos | Endoderm recombination frequency (%) |
| --- | --- | --- | --- |
| R26R<sup>tdTom/tdTom</sup> | Wnt1-Cre2<sup>Tg/+</sup> | 10 | 10 (100) |
| R26R<sup>tdTom/tdTom</sup> | Wnt1-Cre2<sup>Tg/+</sup> ; R26R<sup>tdTom/+</sup> | 23 | 23 (100) |
| Wnt1-Cre2<sup>Tg/Tg</sup> | R26R<sup>tdTom/tdTom</sup> | 26 | 26 (100) |
| Wnt1-Cre2<sup>Tg/Tg</sup> | R26R<sup>eYFP/eYFP</sup> | 3 | 3 (100) |
| Wnt1-Cre2<sup>Tg/Tg</sup> | R26R<sup>mTmG/mTmG</sup> | 3 | 3 (100) |

Anecdotally, the Ai14 R26R^*tdTom*^ reporter line (Madisen et al., 2010) has been activated by cryptic splicing of the STOP cassette upstream of the fluorescent protein. We therefore explored reporter line differences by breeding Wnt1-Cre2 mice with R26R^*eYFP*^ and R26R^*mTmG*^ reporters in a time course of both male and female pups, and in crosses where the Wnt1-Cre2 allele was inherited from either the sire or dam. The Wnt1-Cre2^*Tg/+*^; R26R^*eYFP/+*^ animals exhibited a similar pattern of non-neural crest reporter activity to Wnt1-Cre2^*Tg/+*^; R26R^*tdTom/+*^ mice at postnatal timepoints and into adulthood. However, Wnt1-Cre2^*Tg/+*^; R26R^*mTmG/+*^ animals consistently showed lower penetrance of epithelial signal, which was visually apparent in wholemount [Figure 3A, D]. These differences became clearer in cross-sections made through Wnt1-Cre2^*Tg/+*^; R26R^*eYFP/+*^ and Wnt1-Cre2^*Tg/+*^; R26R^*mTmG/+*^ guts [Figure 3B-C’, E-F’], where the level of recombination in the Wnt1-Cre2^*Tg/+*^; R26R^*mTmG/+*^ was notably less than age-matched Wnt1-Cre2^*Tg/+*^; R26R^*eYFP/+*^ tissue.

**Figure 3:**
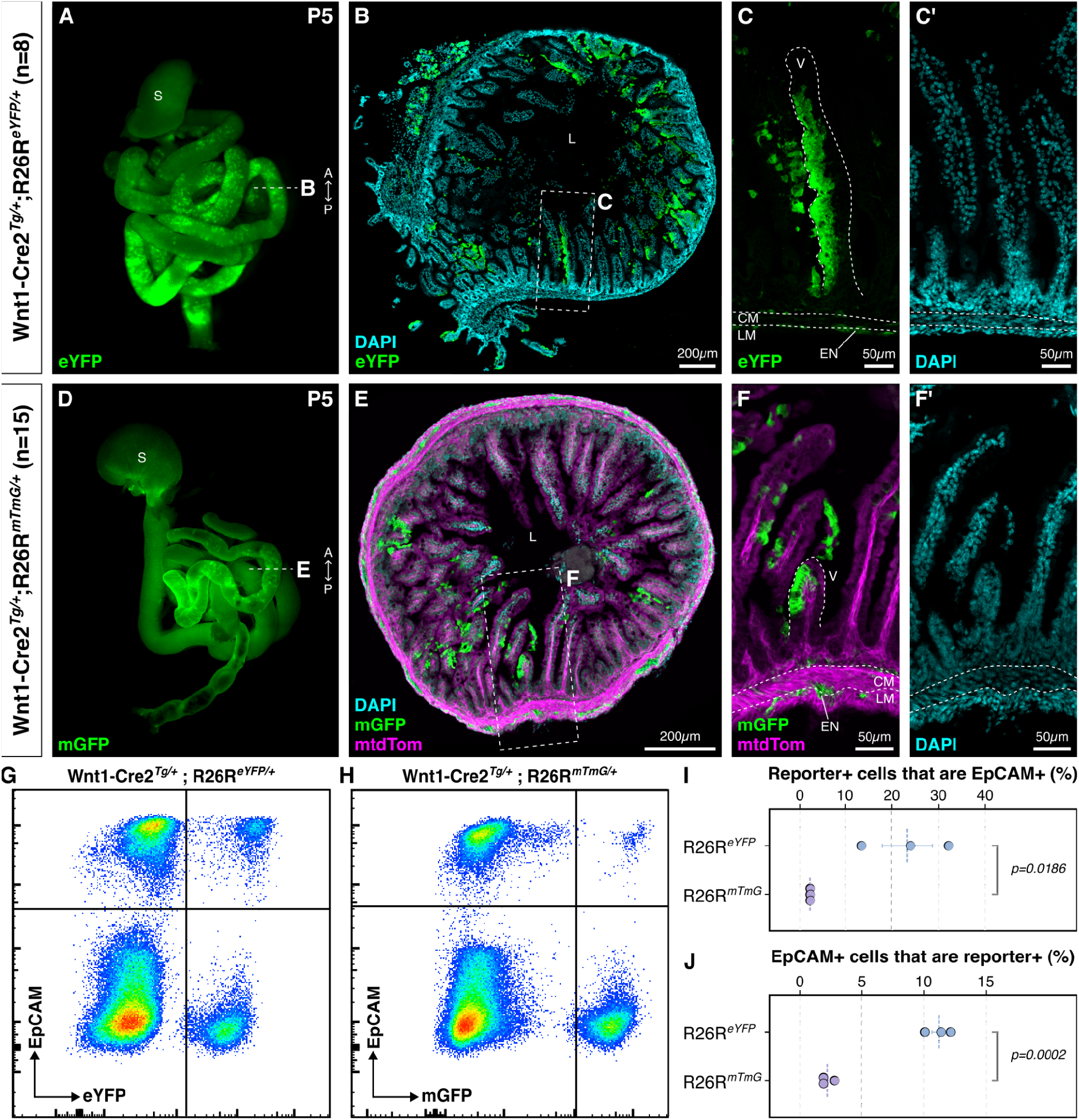
The degree of non-neural crest recombination is reporter-line dependent. **A**. Wholemount gut of a P5 Wnt1Cre2^*Tg/+*^; R26R^*eYFP/+*^ pup. **B**. Transverse cross-section through the small intestine of the tissue shown in A. **C-C’**. Zoomed-in view of reporter activity (C) and nuclei (C’) in the villi. **D**. Wholemount gut of a P5 Wnt1Cre2^*Tg/+*^; R26R^*mTmG/+*^ pup; D-F’ show reduced reporter activation in the gut epithelium when compared to B-C’. **E**. Transverse section through the small intestine of the tissue shown in D. **F-F’**. Zoomed-in view of membrane-GFP reporter activity (F) and nuclei (F’) in the villi. **G-H**. Representative contour plots with biexponential axes of E15.5 Wnt1Cre2^*Tg/+*^; R26R^*eYFP/+*^ (G) and Wnt1Cre2^*Tg/+*^; R26R^*mTmG/+*^ (H) midguts (n=3) with cells exhibiting non-neural-crest reporter activation in the upper-right quadrant. **I**. The proportion of reporter-positive cells that positively stained for EpCAM (n=3 biological replicates, p=0.0186; unpaired two-tailed t-test). **J**. The proportion of EpCAM-positive cells that positively stained for the reporter (n=3 biological replicates, p=0.0002; unpaired two-tailed t-test). (Abbreviations: A, Anterior; P, Posterior; S, Stomach; L, Lumen; V, Villus; CM, Circular Muscle; LM, Longitudinal Muscle; EN, Enteric Neurons)

Reduced penetrance with the R26R^*mTmG*^ reporter line may be due to an increased distance between the loxP sites, where the loxP sites are separated by the coding sequence of the fluorescent reporter and a polyadenylation signal sequence, as opposed to lox-Stop-lox reporters such as R26R^*tdTom*^ and R26R^*eYFP*^, where the proximity of the loxP sites could facilitate their excision in a neural-crest-independent manner. Using flow cytometry, we explored this phenomenon further in midgut tissue dissected from E15.5 Wnt1Cre2^*Tg/+*^; R26R^*eYFP/+*^ and Wnt1Cre2^*Tg/+*^; R26R^*mTmG/+*^ embryos. Using a conjugated antibody against the epithelial cell marker EpCAM, we asked what proportion of epithelial cells in the midgut are reporter-positive, and conversely, what proportion of reporter-positive cells express EpCAM [Figure 3G-H]. Together, we used these statistics as a readout for Cre-mediated off-target recombination.

We observed that 11.2% of EpCAM^+^ cells were eYFP^+^ in Wnt1Cre2^*Tg/+*^; R26R^*eYFP/+*^ embryos, while only 2.20% of EpCAM^+^ cells were mGFP^+^ in Wnt1Cre2^*Tg/+*^; R26R^*mTmG/+*^ embryos [Figure 3I]. This difference was consistent when we looked at the percentage of reporter-positive cells that were EpCAM^+^, with 23.2% of eYFP^+^/EpCAM^+^ cells in Wnt1Cre2^*Tg/+*^; R26R^*eYFP/+*^ embryos, and 2.45% of mGFP^+^/EpCAM^+^ cells in Wnt1Cre2^*Tg/+*^; R26R^*mTmG/+*^ embryos [Figure 3J], suggesting that the choice of reporter can have an impact on the quality of labeling seen with the Wnt1-Cre2 driver line. These results are corroborated by recent work by Drokhlyansky and colleagues (Drokhlyansky et al., 2020), who used the Wnt1-Cre2 driver to label vagal neural crest-derived enteric neurons in the gut for fluorescence-activated cell sorting (FACS) and subsequent transcriptomics. However, they dropped this line from their mouse ENS atlas because of non-specific reporter activation in the colon mucosa, which they traced back to the colon epithelium bioinformatically, even though they used a different fluorescent reporter line (INTACT) compared to the ones we have used in this study. This suggests that the non-neural crest activation of the Wnt1-Cre2 transgene in the C57BL/6J background is independent of the reporter line used.

Wnt1-Cre2 activation has been observed in the male germline, first reported by Dinsmore and Soriano, who proposed that it was a result of Cre activation by regulatory elements controlling the expression of genes surrounding the *E2f1* locus on chromosome 2, in particular, *Actl10, 1700003F12Rik, Necab3*, and *Cbfa2t2* (Dinsmore et al., 2022). If this were true, then we would also expect these genes to influence the expression of Cre recombinase in other cell types where they are expressed. Our histology data suggested that the transgene was active as early as E8.5, when only a subset of the definitive endoderm cells is labeled with the fluorescent reporter. To explore the possibility that genes surrounding the Wnt1-Cre2 locus are driving its expression in the developing mouse gut, we turned to a single-cell RNA-sequencing dataset of splanchnic mesoderm and definitive endoderm cells isolated from E8.5, E8.75, E9.5, and E9.75 embryos (Han et al., 2020).

We reanalyzed the data, subsetting only the annotated definitive endoderm (“de”) cells, further removing 164 cells that were originally labeled as definitive endoderm but clustered together with the splanchnic mesoderm instead. In total, we obtained 4000 cells that split into ten sub-populations [Figure 4B] representing all four (E8.5, E8.75, E9.5, and E9.75) stages [Figure 4C]. All 4000 cells had high levels of *Epcam* expression, confirming their epithelial identity [Figure 4D]. Next, we asked whether genes surrounding the Wnt1-Cre2 locus on the second chromosome are expressed within these cells, focusing on those with coding sequences within 300kb of the *E2f1* gene. We posited that these genes are the most likely to have cis-regulatory elements that could activate Cre in a Wnt1-independent manner. Indeed, eight of the genes analyzed were expressed within the endoderm in varying proportions (Chmp4b - 67%, E2f1 - 23.8%, Cbfa2t2 - 13.5%, Pxmp4 - 9.4%, Zfp341 - 2.3%, Snta1 - 1.7%, and Necab3 - 0.8% of all 4000 cells). One of these genes, *Cbfa2t2*, is a transcriptional co-repressor required for the maintenance of Lgr5+ stem cell populations in the intestine (Amann et al., 2005; Short et al., 2024). These results suggest that one or more enhancers associated with genes flanking the Wnt1-Cre2 transgene can activate Cre expression in non-neural crest lineages, including both male spermatogonia and spermatids (Dinsmore et al., 2022), and the gut epithelium.

**Figure 4:**
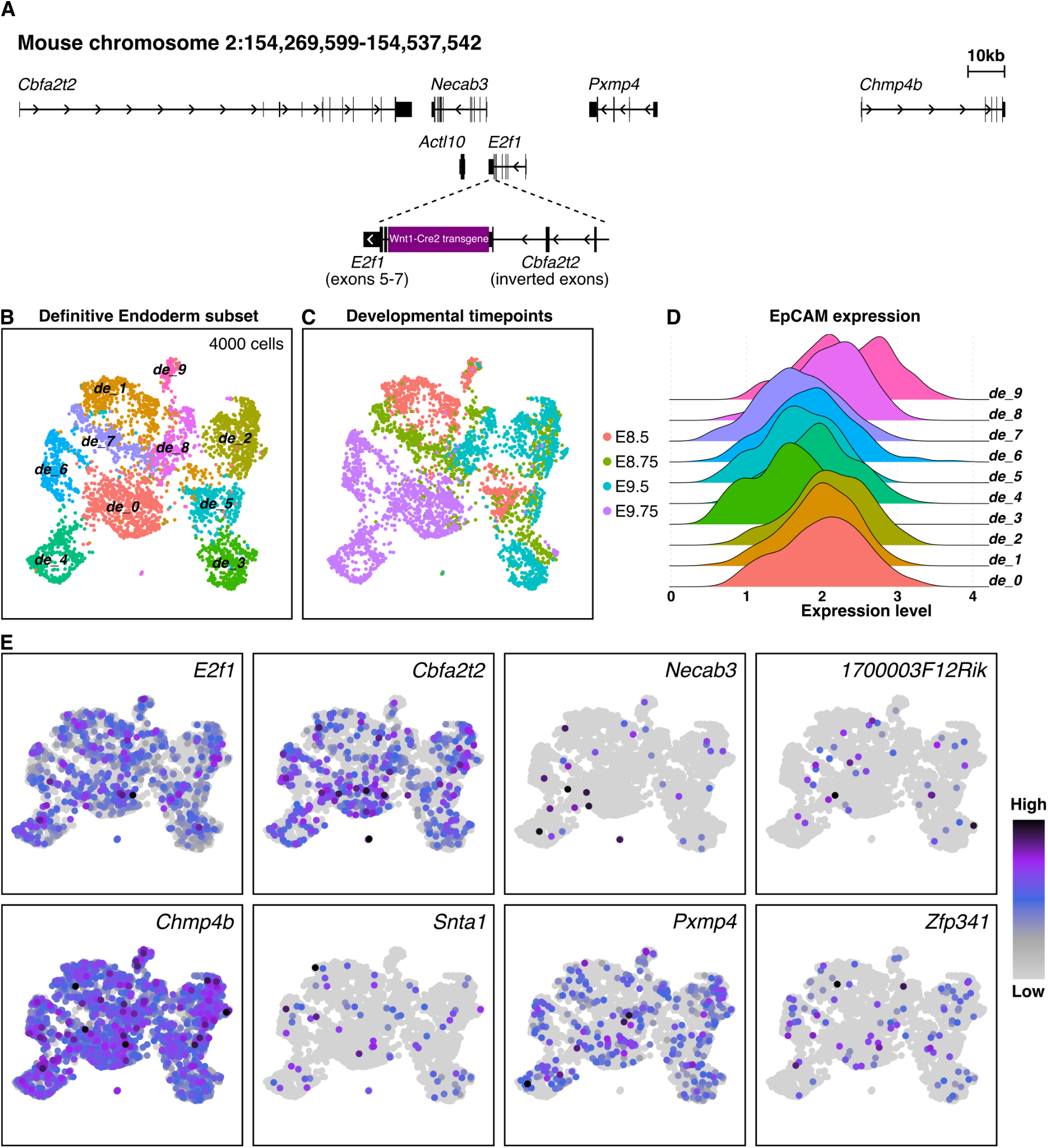
Genes surrounding the Wnt1-Cre2 insertion site are active in the definitive endoderm. **A**. The genomic locus on chromosome 2 of the mouse genome where 1-3 copies of the Wnt1-Cre2 transgene are inserted within the *E2f1* gene. **B**. Reanalysis of the single-cell RNA-sequencing data from definitive endoderm (“de”) cells generated by Han et al., 2020. **C**. The dataset represents four different time points (E8.5, E8.75, E9.5, and E9.75) of mouse embryonic development. **D**. Ridge plot showing the distribution of *Epcam* expression across the ten definitive endoderm sub-clusters. **E**. Feature plots of protein-coding genes with transcription start sites within 300kb of the *E2f1* gene.

In summary, we have demonstrated that the Wnt1-Cre2 driver line maintained on the C57BL/6J background has consistent non-neural crest activation of Cre recombinase, irrespective of the parent. We also show that different reporter lines differ in off-target penetrance when crossed with the Wnt1-Cre2 driver line. In particular, we find that the R26R^*mTmG*^ reporter line has significantly reduced off-target activity of the Cre driver compared to both R26R^*tdTom*^ and R26R^*eYFP*^ lines. However, further work is necessary to identify why and how Cre recombinase is expressed in non-neural-crest cell types. We therefore urge caution with the Wnt1-Cre2 driver for neural-crest-specific lineage tracing and loss-of-function studies and recommend validating phenotypes using other neural crest driver lines.

## Acknowledgments

We thank Kartoosh Heydari, Melaine Delcroix, and Harman Dhaliwal with the Cancer Research Laboratory Flow Cytometry Core Facility at UC Berkeley for assistance with flow cytometry. We thank Ataki Wilson and the rest of the staff at the UC Berkeley OLAC for assistance with rodent breeding strategies and training. We thank Dr. David Drubin and Dr. Megan Martik for allowing access to their confocal microscopes. This work was supported by the NIGMS grant R35GM127069 and C.H. Li Distinguished Professor funds to R.M.H., and a Miller Research Fellowship from the Miller Institute for Basic Research in Science to S.G.

## Conflict of Interest

The authors declare no competing financial or non-financial interests.

## Author contributions

Conceptualization: S.G.; Methodology: S.G., E.J.D.; Validation: S.G., E.J.D.; Formal analysis: S.G., E.J.D.; Investigation: S.G., E.J.D, E.P.; Resources: R.M.H.; Writing - original draft: S.G., E.J.D.; Writing - review & editing: S.G., R.M.H.; Software: S.G.; Visualization: S.G., E.J.D.; Project administration: S.G.; Supervision: S.G., R.M.H.; Funding acquisition: R.M.H.

## Data availability

All mouse lines are available through the Jackson Laboratory.

## Notes

### Competing Interest Statement

The authors have declared no competing interest.

